# Transgenic A53T mice have astrocytic α-synuclein aggregates in dopamine and striatal regions

**DOI:** 10.1101/2025.01.09.632283

**Authors:** Cormac Peat, Asheeta Prasad, David I. Finkelstein, Caroline Sue, Jennifer A. Johnston, Clare L. Parish, Lachlan Thompson, Deniz Kirik, Glenda M. Halliday

**Affiliations:** The University of Sydney, Charles Perkins Centre & Faculty of Medicine and Health School of Medical Sciences, Sydney, NSW, 2050, Australia; Aligning Science Across Parkinson’s (ASAP) Collaborative Research Network, Chevy Chase, MD, 20815, USA; The Florey Institute of Neuroscience and Mental Health & The University of Melbourne, Parkville, VIC, 3052, Australia; Neuroscience Research Australia, Sydney, NSW, 2031, Australia; NysnoBio, Mills Valley, CA, 94941, USA; BRAINS Unit, BMC D11, Lund University, Lund, 22184, Sweden; The University of Sydney, Brain and Mind Centre & Faculty of Medicine and Health School of Medical Sciences, Sydney, NSW, 2050, Australia

**Author notes:** Correspondence to: Professor Glenda Halliday, Brain and Mind Centre, The University of Sydney, Sydney, NSW, 2050, Australia.

**Keywords:** α-synuclein, astrocyte, striatum, substantia nigra, ventral tegmental area

## Abstract

**Aims:** Parkinson’s disease is considered biologically a neuronal α-synuclein disease, largely ignoring the more widespread α-synuclein deposition that occurs in astrocytes, with the aim of this study to identify whether astrocytes accumulate small α-synuclein aggregates before or after neurons.

**Methods:** Fixed serial midbrain and striatal sections from M83 A53T transgenic mouse model of Parkinson’s disease and wild-type controls were histologically processed for multiplex labelling of α-synuclein and astrocytic markers and astrocyte quantitation performed on digital images using QuPath software.

**Results:** The density of astrocytes within the substantia nigra pars compacta was approximately 30% greater compared with other sampled regions (*P*<0.005). Small aggregates of α-synuclein were observed in astrocytic processes, including in wild-type mice where a quarter of all astrocytes had an obvious α-synuclein aggregate. Compared to wild-type, A53T transgenic astrocytes had significantly enlarged somas (*P*<0.001) with more processes (*P*<0.001) consistent with a reactive phenotype. The A53T transgenic mice had more than double the numbers of astrocytes (*P*<0.001) and 2.5 times more astrocytes with α-synuclein aggregates compared to wild-type mice (*P*<0.001).

**Conclusions:** These data suggest that small α-synuclein aggregates are normally cleared by astrocytes and that the substantia nigra pars compacta requires more astrocytic support for this function than other midbrain dopaminergic regions or the striatum. This adds another vulnerability factor to those already known for the substantia nigra with early deficits in clearance of small α-synuclein aggregates by astrocytes associated with an increased astrocytic reactivity in the A53T transgenic mouse model.

**Key Points:**

- The substantia nigra pars compacta contains a higher density of astrocytes than the ventral tegmental area or striatum, indicating a greater reliance on astrocytic function and a greater vulnerability to astrocyte dysfunction
- Small aggregates of α-synuclein were observed in wild-type midbrain and striatal astrocytes, indicating normal clearance of α-synuclein by these astrocytes
- Midbrain and striatal astrocytes from A53T transgenic astrocytes have more than double the number of astrocytes and more astrocytes containing α-synuclein aggregates which have a reactive morphological phenotype.

## Introduction

Parkinson’s disease (PD) is a neurodegenerative disease diagnosed by a loss of dopamine neurons in the substantia nigra pars compacta (SNC) and the widespread accumulation of neuronal intracytoplasmic inclusions of α-synuclein called Lewy pathologies [1]. Genetic mutations in or multiplications of the *SNCA* gene, which encodes α-synuclein, cause familial PD and the first mutation identified in *SNCA* was the A53T mutation [1]. Current concepts of the disease suggest renaming it biologically as neuronal α-synuclein disease [2], but notably, not only do patients with PD deposit α-synuclein in neuritic and neuronal Lewy inclusions, this neuronal pathology is accompanied by α-synuclein deposition in astrocytes [3–5]. In fact, more astrocytes have α-synuclein aggregates (40-50%) than neurons (5-25%) in regions with Lewy pathology in patients with PD [4, 6, 7]. The α-synuclein depositing in astrocytes in PD is non-amyloidogenic, truncated at both the N and C termini for degradation [8, 9] and in isolation (i.e. with limited coexisting neuropathologies) does not cause significant astrocytic reactivity, enlargement or proliferation [7, 10–12]. In fact, there is a negative correlation between increasing α-synuclein accumulation and reduced astrocytic glial fibrillary acidic protein (GFAP) levels in the PD SNC suggestive of a suppressed reactive function [12, 13]}, a reduction in SNC protoplasmic astrocytes [13] and a unique expression of vitamin D-activatin enzyme in PD astrocytes suggestive of an enhanced neuroprotective capability [14]. Understanding the astrocytic biology associated with α-synuclein deposition in PD warrants further analysis, particularly when biologically renaming Lewy body diseases.

Few animal models recapitulate the diagnostic features of PD, but the homozygous A53T α-synuclein transgenic line M83 is the most common model used [15–17]. This model expresses ~5x more human A53T α-synuclein than mouse α-synuclein and develops a delayed severe and complex motor impairment leading to paralysis and death, as well as age-dependent intracytoplasmic neuronal α-synuclein inclusions [15]. Of note, selective high expression (10x increase) of the same A53T transgene in mice astrocytes and not neurons or other brain cells initiates non-cell autonomous killing of neurons in association with the accumulation of aggregated and truncated forms of α-synuclein only in the astrocytes [18]. These mice develop an earlier-onset, rapidly progressive paralysis due to a loss of SNC and spinal motor neurons, as well as a widespread, severe astrogliosis with increased GFAP and excitatory amino acid transporters causing blood-brain-barrier dysfunction and excitotoxicity [18]. This shows that high levels of intracellular α-synuclein activates astrocytes causing considerable dysfunction and surrounding neurodegeneration, something not seen in PD. Until recently, there had been no studies of astrocytic pathology in the most commonly-used M83 mouse model of PD that expresses less of the A53T transgene in all brain cells and has a more slowly progressive and representative form of PD [15–17]. A recent study assessing the blood-brain-barrier in this animal model with neuronal α-synuclein accumulation at 8 months showed decreased expression of tight junction-related proteins, along with increased vascular permeability and accumulation of oligomeric α-synuclein in activated astrocytes in the brain [19], suggesting that intracellular α-synuclein accumulation in astrocytes increases their reactivity and potential to become neurotoxic in these settings [18, 20]. This differs from the lack of astrocytic reactivity with increased α-synuclein accumulation observed in the SNC of PD cases [7, 10–12].

There has been a lot of discussion on the concept and diversity of astrocyte reactivity [21] and this may play a role. Single cell transcriptomics has made it easier to assess astrocytic differences in both human and mice datasets [22]. These comparisons have identified that human astrocytes are much more susceptible to oxidative stress due to lower catalase and glucose-6-phosphate dehydrogenase levels [22]. They have also identified that astrocyte populations in the basal ganglia are similar in both humans and mice and that two subtypes are significantly reduced and two increased in PD [23]. Those sharply decreased in abundance in PD are astrocytes that ensheath neurons (OLIG2+ astrocyte subtype)[24] and reactive A1/inflammation astrocytes (see [25] for subtype), while those increased in abundance are reactive astrocytes involved in oxidase stress and the extracellular matrix (this type is associated with proteostasis in PD) as well as astrocytes restricted to the basal ganglia that are involved in cell-cell and cell-extracellular matrix interactions (LUZP2+ astrocytes)[23]. These changes are similar to the decrease in protoplasmic and increase in fibrous types of astrocytes more recently identified in the PD SNC [13]. Single-cell transcriptomics from the midbrain and striatum of transgenic A53T mice shows an increase in reactive astrocytes with upregulation of nuclear factor kappa-light-chain-enhancer of activation B cells signalling and also vascular endothelial growth factor A (VEGFA)[26], consistent with the recent work on inflammation and blood-brain-barrier disruption in this model [19]. There has been no histological assessment of the abundance of astrocyte types in the basal ganglia of M83 A53T transgenic mice and, while VEGFA+ astrocytes are intimately involved in cell-cell and cell-extracellular matrix interactions in their role regulating the blood-brain-barrier, whether a neuroprotective astrocyte subtype (containing vitamin D-activating enzyme)[14] is increased/observed in this model of PD has not been assessed.

## Materials and methods

### Animals

All animal procedures were conducted in agreement with the Australian National Health and Medical Research Council’s published Code of Practice for the Use of Animals in Research, and approval granted by The Florey Institute of Neuroscience and Mental Health Animal Ethics committee (22-034-FINMH). Brain tissue sections from 6-month-old M83 A53T transgenic mice (*n*=5) and age matched wild type controls (*n*=4) were provided by the Florey Institute of Neuroscience and Mental Health at University of Melbourne and used for analysis. Mice that carry the human A53T mutation driven by the mouse prion promoter (Jax Stock No: 004479; B6;C3-Tg (Prnp-SNCA^*^A53T)83Vle/J; A53T α-synuclein transgenic line M83) were obtained from Jackson Laboratories (resource research identifier (RRID): IMSR_JAX:004479, Bar Harbor, Maine) and a stable breeding colony generated in house (genotype of all mice used in the study confirmed by qPCR)[27]. Mice were euthanised by saline perfusion with 80mg/kg sodium pentobarbital before perfusion with 0.1 M phosphate buffered saline (PBS). Brains were then collected, numbered and left overnight in 4% paraformaldehyde solution before transfer to a 30% sucrose/1M PBS solution, which was refreshed a minimum of two times. Coronal sections were then cut on a cryostat at −25°C and collected serially on 0.5% gelatinised Superfrost slides. Sections were stored desiccated at −80^°^C before the immunohistochemistry was performed.

For the midbrain, 30µm section were sampled every 90µm with three sections mounted per slide giving three sequential series of three sections for each midbrain available for experiments (see below).

For the striatum, 20µm sections were sampled every 200µm with eight sections mounted per slide giving ten sequential series of eight sections from each striatum available for experiments (see below).

### Experimental plan (Table 1)

Experiments were planned to: 1) identify types of astrocytes (using multiple astrocyte markers) and assess whether any particular subtypes of astrocyte contained α-synuclein protein, 2) assess targeted astrocyte markers identified previously as changed in PD (VEGFA [26] and vitamin D-activating enzyme [14]), and 3) quantify the proportion of astrocytes containing α-synuclein protein. Experiments were performed blinded to genotype.

**Table 1:**
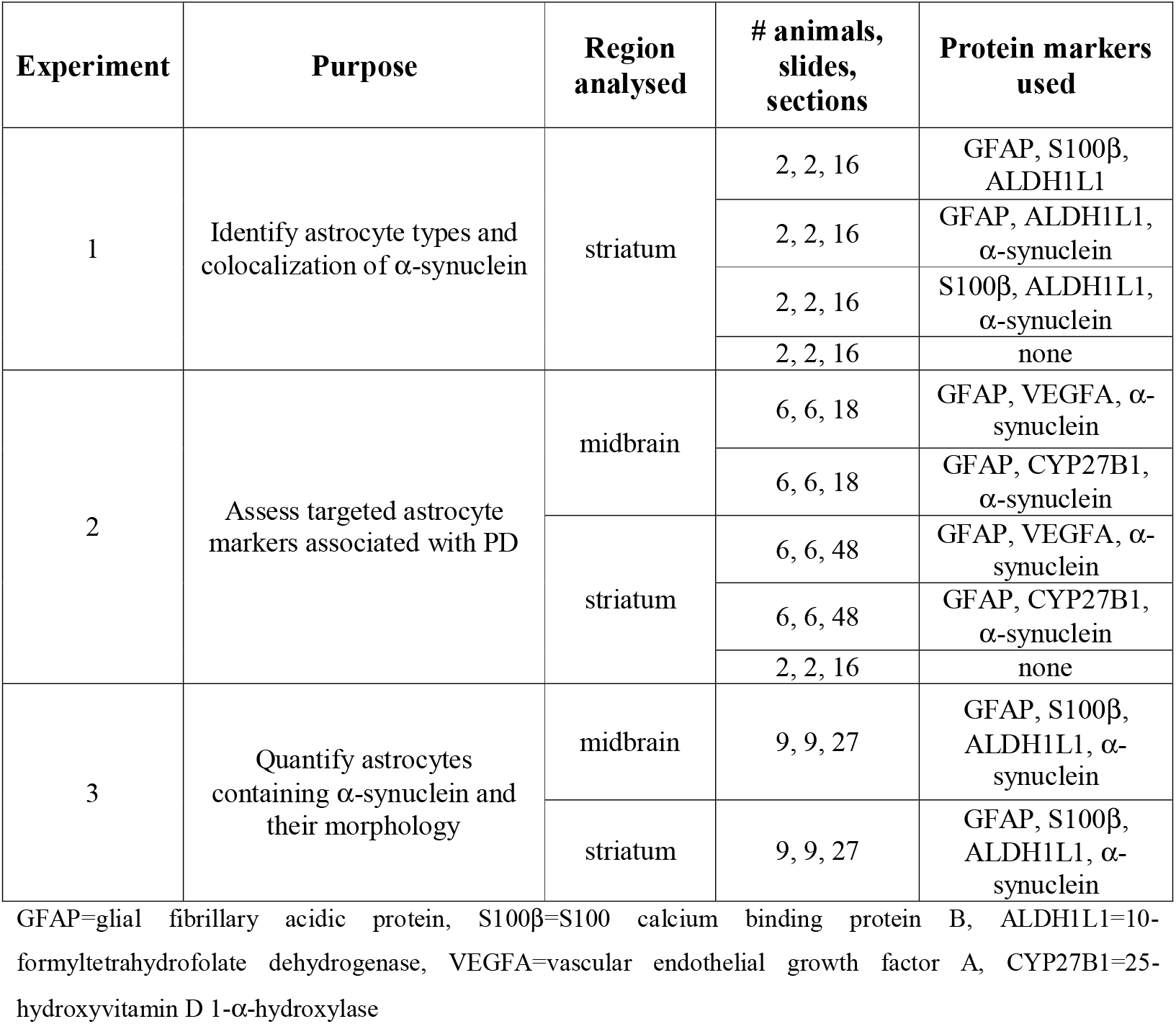
Experimental design.

| Experiment | Purpose | Region analysed | # animals, slides, sections | Protein markers used |
| --- | --- | --- | --- | --- |
| 1 | Identify astrocyte types and colocalization of α-synuclein | striatum | 2, 2, 16 | GFAP, S100β, ALDH1L1 |
| | | | 2, 2, 16 | GFAP, ALDH1L1, α-synuclein |
| | | | 2, 2, 16 | S100β, ALDH1L1, α-synuclein |
|  |  |  | 2, 2, 16 | none |
| 2 | Assess targeted astrocyte markers associated with PD | midbrain | 6, 6, 18 | GFAP, VEGFA, α-synuclein |
| | | | 6, 6, 18 | GFAP, CYP27B1, α-synuclein |
| | | striatum | 6, 6, 48 | GFAP, VEGFA, α-synuclein |
| | | | 6, 6, 48 | GFAP, CYP27B1, α-synuclein |
|  |  |  | 2, 2, 16 | none |
| 3 | Quantify astrocytes containing α-synuclein and their morphology | midbrain | 9, 9, 27 | GFAP, S100β, ALDH1L1, α-synuclein |
| | | striatum | 9, 9, 27 | GFAP, S100β, ALDH1L1, α-synuclein |

GFAP=glial fibrillary acidic protein, S100β =S100 calcium binding protein B, ALDH1L1=10-formyltetrahydrofolate dehydrogenase, VEGFA=vascular endothelial growth factor A, CYP27B1=25-hydroxyvitamin D 1- $\alpha$ -hydroxylase

### Immunofluorescence

Briefly, sections were washed in 1X tris-buffered saline (TBS) with 1.25% Triton X-100 and blocked for one hour in bovine serum albumin (BSA)/casein blocking solution (4% BSA (w/v), 1% casein (w/v), 1.5% glycine (w/v), 0.25% Triton X-100 (v/v) in 1xPBS), then incubated with a combination of primary antibodies overnight at 4°C (see Tables 1 and 2). After washing with PBS, the tissue sections were incubated with appropriate fluorophore-conjugated secondary antibodies for three hours (see Table 2), then thoroughly washed with TBS, counterstained with Hoechst (Thermo Fischer Scientific Cat# 62249; 1ug/mL) before a final wash and mounting with Prolong Diamond Antifade mounting medium (Invitrogen Cat# P36970), then coverslipped. Control experiments with incubation without primary antibodies gave no immunofluorescence signals.

**Table 2:** Primary and secondary antibodies used.

| Marker | Primary antibodies | Secondary antibodies |
| --- | --- | --- |
| GFAP | chicken anti-GFAP (Thermo Fisher Scientific Cat# PA1-10004, RRID:AB_1074620; diluted 1:1000) | Goat anti-chicken 488 (Thermo Fisher Scientific Cat# A-11039, RRID:AB_2534096; diluted 1:250) or Goat anti-chicken 594 (Thermo Fisher Scientific Cat# A-11042, RRID:AB_2534099; diluted 1:250) or Goat anti-chicken 647 (Thermo Fisher Scientific Cat# A-21449, RRID:AB_2535866; diluted 1:250) |
| S100β | goat anti-S100β (R and D Systems Cat# AF1820, RRID:AB_2184423; diluted 1:1000) | Donkey anti-goat 647 (Thermo Fisher Scientific Cat# A-21447, RRID:AB_2535864; diluted 1:250) or Donkey anti-goat 594 (Thermo Fisher Scientific Cat# A-11058, RRID:AB_2534105; diluted 1:250) |
| ALDH1L1 | rabbit anti-ALDH1L1 (Thermo Fisher Scientific Cat# PA5-78622, RRID:AB_2735443; diluted 1:1000) | Goat anti-rabbit 647 (Thermo Fisher Scientific Cat# A32733, RRID:AB_2633282; diluted 1:250) or Goat anti-rabbit 594 (Thermo Fisher Scientific Cat# A-11012, RRID:AB_2534079; diluted 1:250) |
| α-synuclein | Mouse anti-alpha-synuclein (BD Biosciences Cat# 610787, RRID:AB_398108; diluted 1:1000) | Goat anti-mouse 488 (Thermo Fisher Scientific Cat# A-11001, RRID:AB_2534069; diluted 1:250) |
| VEGFA | Rabbit anti-VEGF (GeneTex Cat# GTX102643, RRID:AB_11174248; diluted 1000) | Goat anti-rabbit 594 (Thermo Fisher Scientific Cat# A-11012, RRID:AB_2534079; diluted 1:250) |
| CYP27B1 | Rabbit anti-CYP27B1 (Millipore Cat# ABN182, RRID:AB_2616605; diluted 1:1000) | Goat anti-rabbit 594 (Thermo Fisher Scientific Cat# A-11012, RRID:AB_2534079; diluted 1:250) |

GFAP=glial fibrillary acidic protein, S100β =S100 calcium binding protein B, ALDH1L1=10-formyltetrahydrofolate dehydrogenase, VEGFA=vascular endothelial growth factor A, CYP27B1=25-hydroxyvitamin D 1- $\alpha$ -hydroxylase

For all experiments (Table 1), triple labelling immunofluorescence with non-overlapping flourophores (Table 2) were used to determine colocalization. For experiment 3 (Table 1), GFAP, S100 calcium binding protein B (S100β) and aldehyde dehydrogenase 1 family member L1 (ALDH1L1) were all identified by the same flourophore (Table 2) to give the maximum labelling of astrocytes for colocalization with α-synuclein and quantitation (nuclei were counterstained with Hoechst).

### Image capture, processing and analysis

Sections were visualised at 10x magnification with a Leica THUNDER 3D Imager and standardised regions 200 x 200µm in size selected. Z stack images of these regions were captured at 1μm intervals for 10μm at 40X/NA1.1 magnification (this z distance separated overlapping astrocytes), and an additional high magnification image (63X/NA1.4) taken to ensure resolution of intracellular astrocytic labelling and for morphological measurements. The brightness and contrast of the images was standardised. For the midbrain slide, one region in the SNC and one region in the ventral tegmental area (VTA) was captured at the three levels in the series (Fig. 1). For the larger striatum, standardised rostral, mid and caudal levels were selected, and four regions captured per level in the series (Fig. 1).

**Figure 1.**
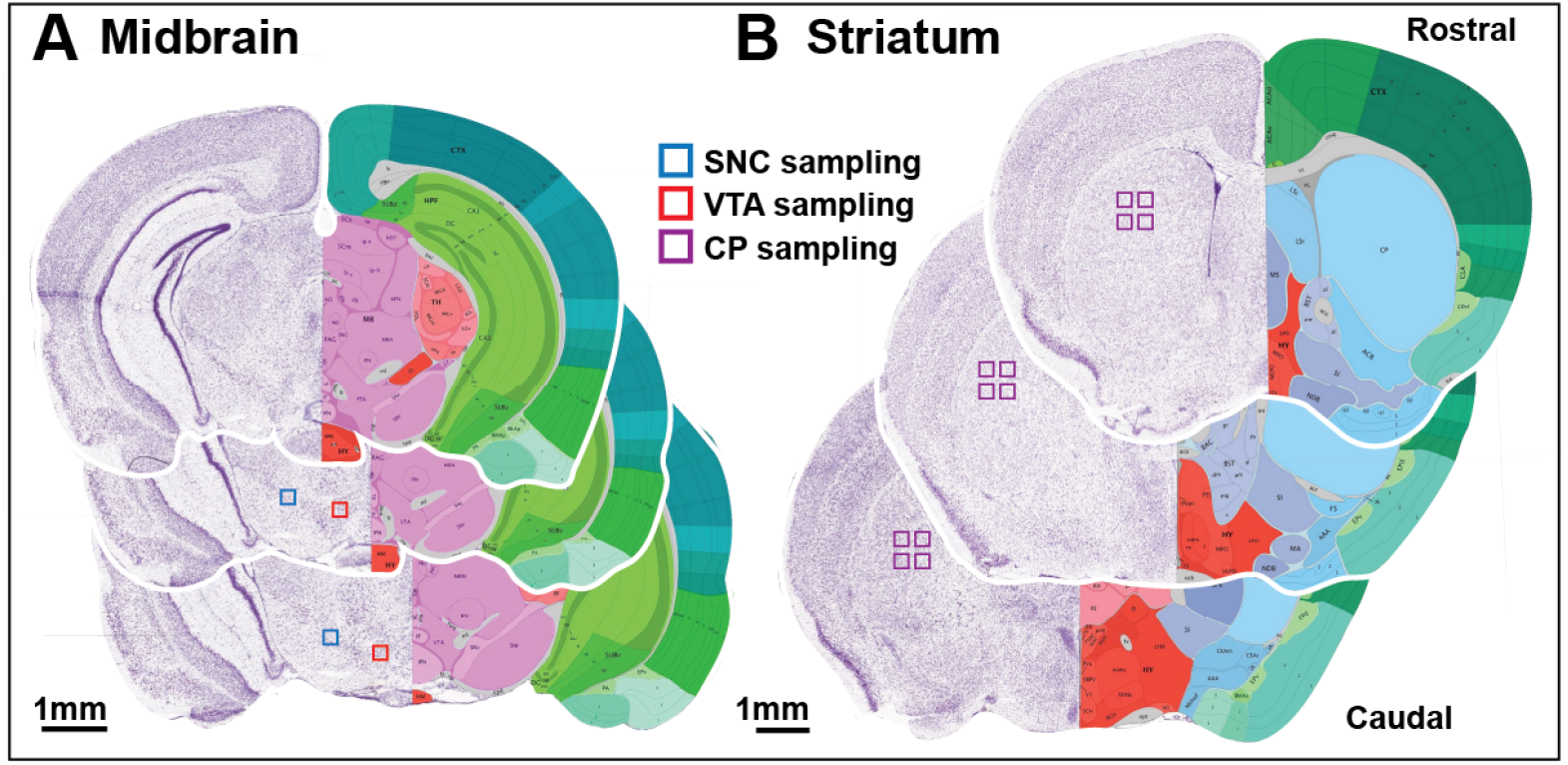
Sampling schematic for image capture and cell quantification in 200×200µm sampling boxes. **(A)** For each of the three midbrain levels spaced 90µm apart (anterior at top), a single box (blue) was sampled in the substantia nigra pars compacta (SNC) and another (red) in the ventral tegmental area (VTA) giving 3 sampling boxes per region. **(B)** For each of the three striatal (abbreviated CP) levels spaced 600µm apart (anterior at top), four boxes (purple) were sampled at each level giving a total of 12 sampling boxes. Images are from standardised coronal Allen Adult Mouse Brain Atlas (https://mouse.brain-map.org/experiment/thumbnails/100048576?image_type=atlas). All data is standardised to the sampling box area for each region of interest.

For experiment 3, 10-15 astrocytes displaying an obvious soma and clear processes were first established by separating the image by channel (AlexaFluor 594 for all astrocyte markers, experiment 3 in Table 1) in the high magnification images (63X/NA1.4) taken in each region of interest (ROI). These representative astrocytes were assessed for any changes in astrocyte morphology (soma size and the number of astrocyte processes). The soma of each astrocyte was traced using the polygon annotation tool in QuPath and the area measured. The number of processes for each astrocyte was also recorded.

In the entire ROI images, the widefield fluorescence Z stack images were exported as tiffs, processed and flattened using the QuPath software (Bankhead et al., 2017) for quantitation. The total number of astrocytes was established by separating the image by channel (AlexaFluor 594 for all astrocyte markers, experiment 2 in Table 1) and counting all astrocytes identified in the Z stack manually. Only astrocytes that displayed an obvious soma and clear processes were included. Once the total number of astrocytes were counted, double labelled astrocytes (astrocyte markers and α-synuclein, see Table 1) were counted manually and the percentage of astrocytes expressing α-synuclein determined by dividing the number of double-labelled astrocytes by the total number of astrocytes.

Images were colour changed to ensure colour-blind compatibility for presentation. Immunofluorescence for triple labelled GFAP+S100β+ALDH1L1 or for single labelled GFAP was changed to cyan, for single labelled S100β or α-synuclein was changed to magenta, for single labelled ALDH1L1, VEGFA or for the vitamin D activator CYP27B1 was changed to yellow, and for DAPI was changed to grey.

### Statistical analysis

Statistical analyses were performed using IBM SPSS Statistics (Version 29.0.2.0(20)) and data is presented as mean ± SEM. *P* < 0.05 was taken as statistically significant.

To assess the effects of genotype on regional astrocyte morphology and numbers, multivariate linear regression for the soma size of astrocytes and the number of astrocyte processes (genotype by region), as well as for the total astrocyte numbers and the proportion of astrocytes with α-synuclein aggregates (genotype, region and level (see Fig. 1)) was used with Tukey post-hoc tests.

## Results

### Types of astrocytes observed in the midbrain and striatum

Astrocyte subtypes were analysed in the striatum where subtypes have been previously identified [23] as well as two regions of the midbrain. The different pan astrocyte markers used (Table 2) identified the same cells with similar morphologies, but the protein localisation for GFAP (astrocytic cytoskeletal marker), S100β (astrocytic calcium binding protein) and ALDH1L1 (astrocytic mitochondrial metabolising enzyme) within each astrocyte differed (Fig. 2). In WT mice, GFAP was observed throughout the soma and processes of most astrocytes (less soma and more processes identified in controls, not shown), while ALDH1L1 and S100β had lower fluorescence within the processes of some astrocytes (Fig. 2). Overall, the different markers did not label different astrocyte subtypes in the striatum or midbrain and therefore further experiments used either GFAP (which was observed in both the soma and processes of many astrocytes) or all three markers (to maximise the number of astrocytes observed)(Table 1).

**Figure 2.**
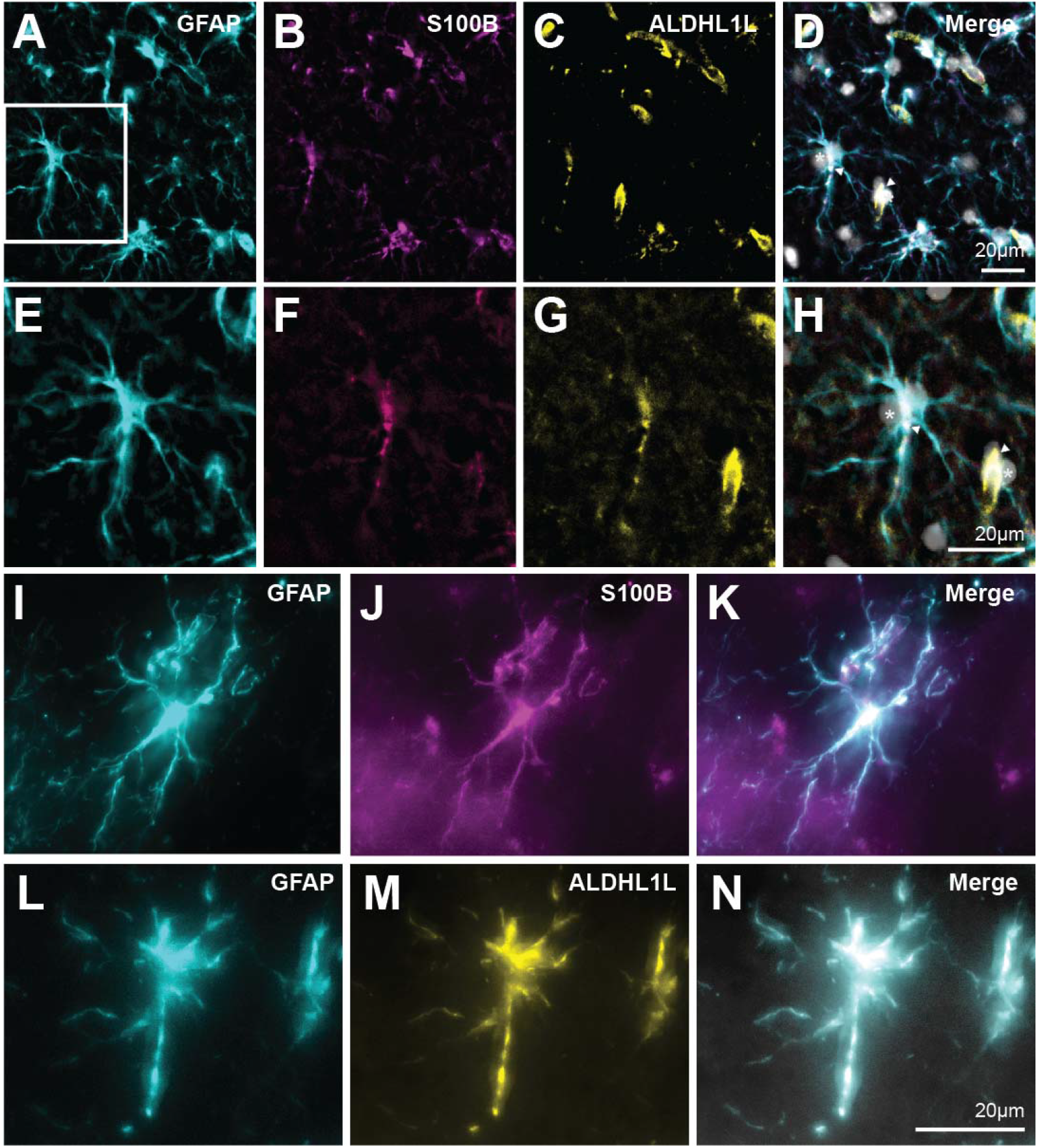
Evaluation of astrocyte markers for diverse astrocyte types. **(A-H)** Triple labelled immunofluorescence in the striatum of wild-type mice using antibodies to the cytoskeletal protein GFAP (cyan in A and E), the calcium binding protein S100β (magenta in B and F) and the astrocytic mitochondrial metabolising enzyme ALDH1L1 (yellow in C and G). E-H are the enlarged region indicated in A. Scale in D is equivalent for A-C. Scale in H is equivalent for E-G. These proteins were found co-localised in the same astrocytes in the striatum (overlap white in D and H and arrowed, DAPI is grey and some nuclei asterisked). **(I-K)** Double labelled immunofluorescence of a striatal astrocyte showing GFAP (cyan in I) and S100β (magenta in J) co-localisation (white in K). **(L-N)** Double labelled immunofluorescence of midbrain astrocytes showing GFAP (cyan in L) and ALDH1L1 (yellow in M) co-localisation (white in N). Scale in N equivalent for I-M.

### Most astrocytes in A53T transgenic mice appear reactive

The morphology of the astrocytes in all regions examined in the A53T transgenic mice were consistent with reactive astrocytes where they upregulate core markers including the astrocytic cytoskeletal marker GFAP, the astrocytic calcium binding protein S100B, and the astrocytic mitochondrial metabolising enzyme ALDH1L1 [21], all the markers used in the present study. Astrocytes in the A53T transgenic mice had significantly enlarged somas (Figs. 3,4A-G, mean µm^2^±SEM A53T=0.458±0.035, WT=0.216±0.039, *F*(1)=21, *P*<0.001) across all regions (*F*(2)=0.32, *P*=0.728) and an increased number of processes (Figs. 3,4A-F,H, mean±SEM A53T=4.19±0.14, WT=3.10±0.16, *F*(1)=27, *P*<0.001) across all regions (*F*(2)=0.17, *P*=0.849) compared with WT mice.

**Figure 3.**
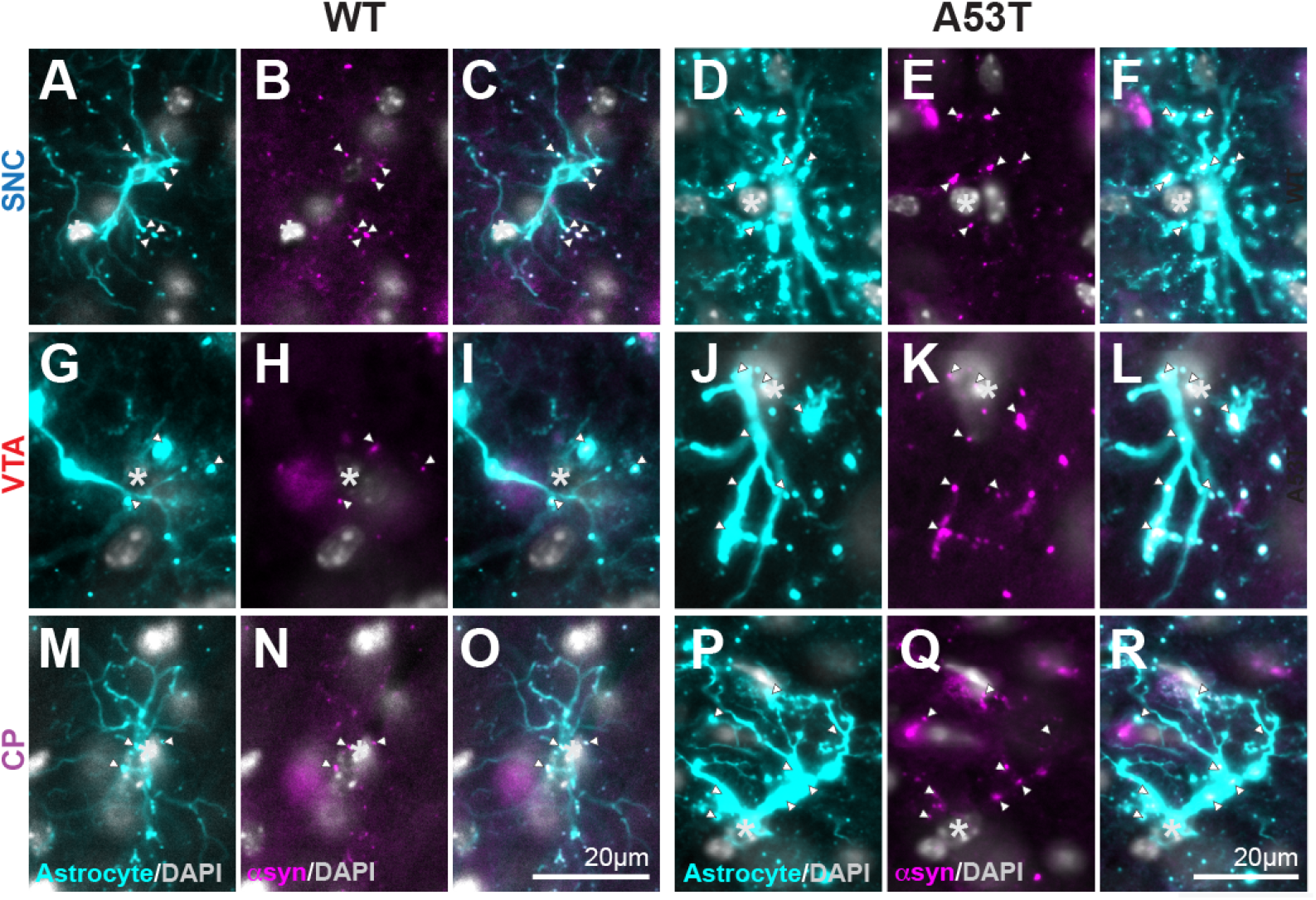
α-Synuclein puncta are common in astrocytic processes. Immunofluorescence labelling of astrocytes using all three astrocytic proteins (GFAP, S100ß and ALDH1L1 cyan) and α-synuclein (magenta) in the substantia nigra pars compacta (SNC, A-F), ventral tegmental area (VTA, G-L) and striatum (CP, M-R) of wild-type (WT, A-C, G-I, M-O) and A53T transgenic (A53T, D-F, J-L, P-R) mice showing α-synuclein aggregates in their processes (white arrowheads). Grey=nuclear DAPI. Scale bar in O is equivalent for WT micrograophs (A-C, G-I, M-O). Scale bar in R is equivalent for A53T micrographs (D-F, J-L, P-R).

**Figure 4.**
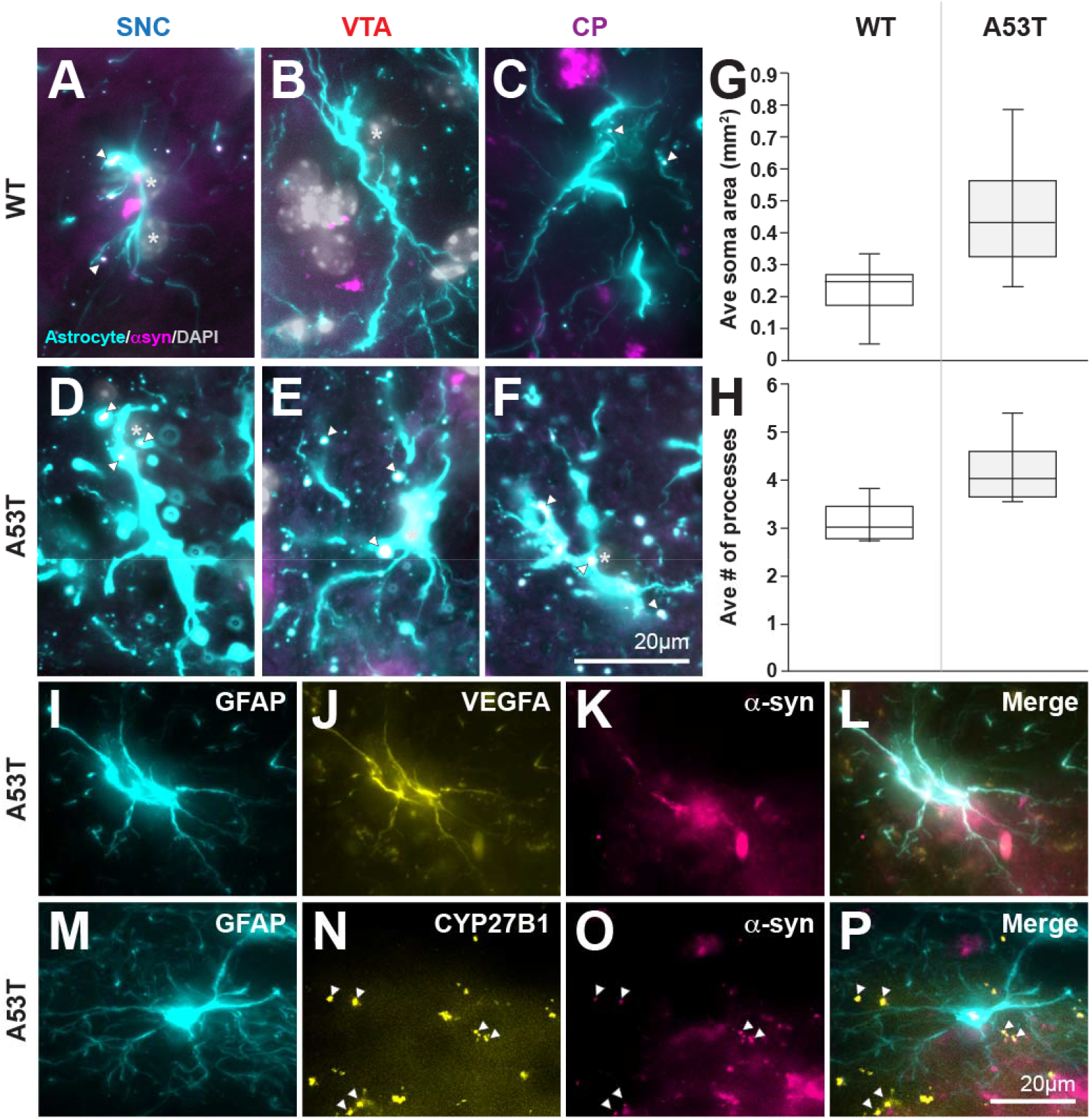
Evidence of reactive astrocytes in A53T transgenic mice. **(A-F)** Immunofluorescence labelling of astrocytes using all three astrocytic proteins (GFAP, S100ß and ALDH1L1 cyan) and α-synuclein (magenta) in the substantia nigra pars compacta (SNC, A,D), ventral tegmental area (VTA, B,E) and striatum (CP, C,F) of wild-type (WT, A-C) and A53T transgenic (A53T, D-F) mice showing α-synuclein aggregates in their processes (white arrowheads). Grey=nuclear DAPI. Scale bar in F is equivalent for A-E **(G)** Boxplot of the median astrocyte soma area and the interquartile range (in µm^2^) in wild-type (WT, *n*=4) and A53T transgenic (A53T, *n*=5) mice showing more than a doubling in the soma area of astrocytes in A53T versus WT mice across all regions assessed using multivariate linear regression (*F*_soma size_(1)=21, *P*<0.001; *F*_region_(2)=0.32, *P*=0.728) consistent with a reactive phenotype. **(H)** Boxplot of the median number of processes per astrocyte and the interquartile range in wild-type (WT, *n*=4) and A53T transgenic (A53T, *n*=5) mice showing a 35% increase in the number of astrocytic processes in A53T versus WT mice across all regions assessed using multivariate linear regression (*F*_processes_(1)=27, *P*<0.001; *F*_region_(2)=0.17, *P*=0.849) consistent with a reactive phenotype. **(I-L)** Triple labelled immunofluorescence in the striatum of A53T transgenic (A53T) mice using antibodies to the cytoskeletal protein GFAP (cyan in I), the growth factor for vasculature and blood-brain-barrier VEGFA (yellow in J) and α-synuclein (magenta in K) showing that reactive astrocytes in A53T mice with α-synuclein aggregates express VEGFA (white in L) consistent with increased reactivity as recently identified [19]. **(M-P)** Triple labelled immunofluorescence in the striatum of A53T transgenic (A53T) mice using antibodies to the cytoskeletal protein GFAP (cyan in I), the synthesising enzyme for vitamin D CYP27B1 (yellow in J) recently identified throughout the cytoplasm of astrocytes that sequester α-synuclein in sporadic Parkinson’s disease [14], and α-synuclein (magenta in K) showing that reactive astrocytes in A53T mice with α-synuclein aggregates do not express CYP27B1 throughout their cytoplasm, but can have rare vesicular-like aggregates (white arrowheads).

Assessment of example functional markers also indicated that the reactive astrocytes in A53T transgenic mice had functional changes. Vascular coupling and blood-brain-barrier integrity was assessed with VEGFA which confirmed that most reactive astrocytes in the A53T transgenic mice were also VEGFA+ (Fig. 4I-K), as recently published [19]. In contrast to the specific expression of the synthesising enzyme for vitamin D CYP27B1 in the entire cytoplasm in astrocytes that sequester α-synuclein in the brains of sporadic PD patients [14], this PD-specific change was not observed in A53T transgenic mice (Fig. 4M-P). Only small vesicles containing both CYP27B1 and α-synuclein were found in A53T transgenic mice astrocytes with α-synuclein aggregates (Fig. 4M-P).

### Most astrocytes in A53T transgenic mice accumulate α-synuclein

Assessment of astrocytes combining all astrocyte marker proteins together allowed their full morphology to be identified (Figs. 3,4A-F) and assessment of α-synuclein aggregates within them easier to identify. Small aggregates of α-synuclein were observed mostly in astrocytic processes, even in WT mice (Figs. 3,4A, arrowheads) where just over one in four astrocytes had an obvious α-synuclein aggregate in one or more processes (Fig. 5B). This suggests that aggregates of α-synuclein are normally cleared by astrocytes.

**Figure 5.**
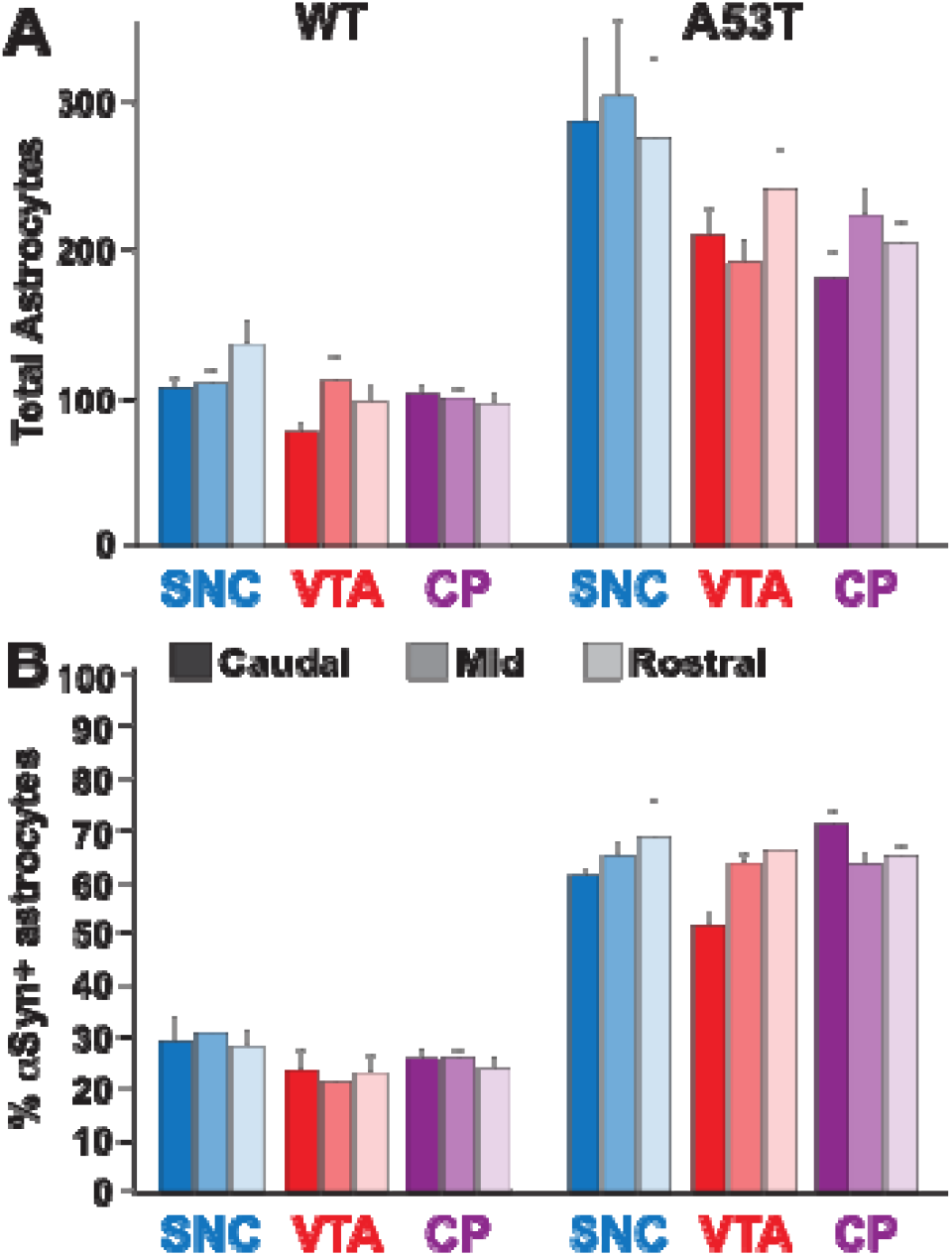
Evidence for a significant increase in the number of astrocytes in A53T transgenic mice. **(A)** Graph of the total number of astrocytes per regional sample box by genotype (wild-type WT *n*=4 and A53T transgenic *n*=5), region (substantia nigra pars compacta SNC, ventral tegmental area VTA and striatum CP) and regional level (rostral, middle and caudal, see Figure 1) showing no significant differences between the density of astrocytes at different levels of the same regions using multivariate linear regression (*F*_location_(2)=0.33, *P*=0.72). Multivariate linear regression revealed that the SNC had an increased density of astrocytes (approximately 30% more) compared with the VTA and CP (*F*_region_(2)=6.8, *P*=0.002, posthoc tests between SNC and other regions *P*<0.005) and that A53T mice had more than double the average number of astrocytes per sample box compared with WT mice (*F*_genotype_(1)=104, *P*<0.001) with no genotype by region interactions (*F*(2)=3.085, *P*=0.053). **(B)** Graph of the proportion of astrocytes containing α-synuclein aggregates by genotype (wild-type WT *n*=4 and A53T transgenic *n*=5), region (substantia nigra pars compacta SNC, ventral tegmental area VTA and striatum CP) and regional level (rostral, middle and caudal, see Figure 1) showing no significant differences between proportion of α-synuclein-containing astrocytes at different levels of the same regions using multivariate linear regression (*F*(2)=0.077, *P*=0.93). Multivariate linear regression revealed that the VTA had less α-synuclein-containing astrocytes (just over 10% less) compared with the SNC (*F*_region_(2)=3.158, *P*=0.049, posthoc tests between SNC and VTA *P*<0.02) and that A53T transgenic mice had nearly a 2.5x greater proportion of astrocytes with α-synuclein aggregates compared to WT mice (*F*_genotype_(1)=494, *P*<0.001) with no genotype by region interactions (*F*(2)=0.905, *P*=0.41).

Multivariate analysis revealed a significant difference in the density of astrocytes and the proportion of astrocytes containing α-synuclein aggregates (Fig. 5) between genotype (*F*(2)=420, *P*<0.001) and region (*F*(4)=5.95, *P* <0.001) but not level within region (*F*(4)=0.291, *P*=0.88). There was no genotype by region interactions (*F*(2,4)=1.54, *P*=0.20). In A53T transgenic mice, the majority of astrocytes contained α-synuclein aggregates (Fig. 5B), and compared with WT astrocytes, the α-synuclein aggregates were observed in more than one process and often also within their soma (Figs. 3,4D-F). Diffuse α-synuclein accumulations in occasional neurons was also seen in each of the areas analysed (data not shown), although frank Lewy pathology was not observed. The A53T transgenic mice had more than double the average number of astrocytes per sample box compared with WT mice (mean±SEM A53T=249±10, WT=104±11, *F*(1)=104, *P*<0.001, Fig. 5A). A53T transgenic mice also had nearly a 2.5x greater proportion of astrocytes with α-synuclein aggregates compared to WT mice (mean±SEM A53T=64±1%, WT=26±1%, *F*(1)=494, *P*<0.001, Fig. 5B). There were approximately 30% more astrocytes in the SNC compared with the VTA and striatum (CP) (mean±SEM SNC=213±12, VTA=162±12, CP=153±12, *F*(2)=6.8, *P*=0.002, posthoc *P*<0.005, Fig. 5A) suggesting this region requires more astrocytic support.

## Discussion

PD is considered to be a neuronal α-synuclein disease [2], but the main neuronal pathologies are accompanied by significantly more α-synuclein deposition in astrocytes which is poorly understood [3–5, 7]. The astrocytes with α-synuclein accumulation in sporadic PD do not have a reactive phenotype [7, 10–12] (unless Alzheimer co-pathology occurs)[28, 29] and have neuroprotective properties, like expressing the vitamin D-activating enzyme particularly around neurons without Lewy pathologies [14]. Cell studies have identified that astrocytes can ingest enormous amounts of α-synuclein of different forms and are activated to degrade the α-synuclein [30–33]. In these models, astrocytes can store α-synuclein due to incomplete lysosomal degradation, which with progressive accumulation can cause mitochondrial impairments in the astrocytes and senescence, affecting their function [31, 34]. This is likely to be relevant for PD as this important astrocyte function can be induced by PD-linked environmental factors [35]. These studies suggest that effective astrocyte function is essential for adequate removal of pathogenic α-synuclein and that factors impacting on this function increase α-synuclein accumulation in the brain.

The normal accumulation of the prevalent protein α-synuclein in astrocytes has not been identified in the brains of WT mice previously, but the careful examination of astrocytes in the present study revealed α-synuclein accumulations in the processes of astrocytes (Figs. 3,4A, C). Just over one in four WT astrocytes had α-synuclein aggregates in their processes (Fig. 5B). Of note, no α-synuclein accumulations were observed in WT neurons in these regions, suggesting that astrocytes normally remove any excess α-synuclein. This has only been observed previously in *in vitro* experiments where similar isolated α-synuclein puncta are found in around 40% of astrocytes after 24 hours [31] with phagocytic accumulation of α-synuclein peaking at 48 hours and the α-synuclein degraded by 170 hours [30]. Interestingly, the SNC has approximately 30% more astrocytes than the VTA or CP in mice (Fig. 5A) suggesting that more astrocyte support is required in this region. In the SNC the density of neurons is significantly increased as the name suggests (compact) and in humans a single astrocyte is associated with each dopamine neuron in the SNC whereas in the CP a single astrocyte is associated with six to eight neurons [36]. The increased astrocytes in the SNC in mice could indicate a similar working arrangement in the SNC of mice where a single astrocyte cares for each dopamine neuron. Together these data confirm the importance of astrocytes in disposing of excess α-synuclein in normal healthy brain tissue and that this relationship appears most intimate in the SNC. While speculative, this process may be related to astrocyte-dependent circuit remodelling by synapse phagocytosis that is essential for synaptic homeostasis [37]. This is a process that has not been well studied in these regions in mice but astrocytic synaptic and dendritic spine remodelling are known to occur in progressive models of PD [38, 39].

Animal models of PD have not assessed astrocytic accumulation of α-synuclein to any extent. In the present study assessment of astrocytes was performed at 6 months prior to the onset of motor dysfunction and dopamine neuron loss, which begin between 8-12 months of age in this model. Accumulation of oligomeric α-synuclein in activated astrocytes in the brain of 8 month old A53T transgenic mice was reported recently in association with decreased expression of tight junction-related proteins and increased vascular permeability [19]. As this study focussed on blood-brain-barrier disruption in the symptomatic stage, no general quantitation of the impact of any α-synuclein accumulation on astrocytes was evaluated. The present study quantitatively assessed the reactivity of astrocytes prior to symptom onset and found significant accumulation of α-synuclein in association with increased astrocyte reactivity in A53T transgenic mice at 6 months. The astrocytes in this common PD model did not have dispersed α-synuclein accumulation throughout their cytoplasm at this age, as often seen in end-stage PD in areas with significant Lewy pathology [3–5, 7], but they had an increase in the presence of α-synuclein aggregates in their processes as well as in their somas (Figs. 3,4D-F). In fact, most astrocytes contained α-synuclein aggregates in this PD mouse model at this early stage (Fig. 5B). While the astrocytes in the A53T transgenic mice were more reactive (twice as many that were double the size with more processes (Figs. 4G,H,5A)), their increased size appeared commensurate with the increase in the amount of α-synuclein aggregates in each astrocyte, suggestive of an increase in the turnover of these aggregates by these more reactive astrocytes. Similar morphological changes in perisynaptic astrocytes have been described in the rat CP following 6-OHDA dopaminergic denervation [40, 41]. At the age assessed (6 months), the A53T transgenic mice had diffuse α-synuclein accumulations in some neurons in each of the areas analysed (data not shown), although frank Lewy pathology was not observed. This indicates that the increased astrocytic α-synuclein occurs before significant neuronal α-synuclein begins in this model. This is also observed in the preformed fibril mouse model with injection into the CP where astrocyte reactivity occurs significantly earlier that α-synuclein inclusion formation in SNC dopaminergic neurons and is associated with striatal dopamine terminal denervation [42].

As recently detailed at 8 months [19], the reactive astrocytes in the A53T transgenic mice also contained VEGFA at 6 months (before symptom onset), an indicator of functional upregulation of vascular coupling, increased hypoxia sensing and reduced blood-brain-barrier integrity. Reduced blood-brain-barrier integrity and an increase in striatal blood vessel density has been shown in human α-synuclein overexpressing mice at 8 months, followed at 13 months (now symptomatic) by more significant accumulation of α-synuclein, a decrease in dopamine terminals and a decrease in blood vessel density [43]. These data suggest that blood-brain-barrier disruption occurs with increasing astrocytic VEGFA and α-synuclein accumulation in astrocytes. While there is limited data on VEGFA in the PD brain, elevated immunoreactivities in the SNC but not striatal astrocytes with increased GFAP has been reported [19, 44] with an increase in vascular endothelial cells [45] suggesting some neovascularisation. In the adult rat rotenone model of PD, the SNC also has an increase in the number of astrocytes and blood capillaries due to an increase in string vessels [46]. SNC string vessel formation has been identified in PD brain with increased astrocytes in the striatum of the same cases reducing this effect in this location [47]. The data suggest that astrocyte function supporting blood vessels in the SNC in particular is disrupted with increasing astrocytic α-synuclein accumulation.

As indicated, the highly reactive astrocytes observed in the PD model studied do not occur in the PD SNC [7, 10–12], which may be a function of the disease stage studied (early in the present study, later in most human tissue studies). In humans with sporadic PD there is a neuroprotective upregulation of the synthesising enzyme for vitamin D CYP27B1 throughout the cytoplasm of the 50% of astrocytes that contain α-synuclein puncta and are associated with neurons that do not contain Lewy pathology [14]. However, the mice astrocytes in the present study did not have widespread astrocytic cytoplasmic CYP27B1 that indicates a neuroprotective phenotype, but did have restricted co-localisation of CYP27B1 and α-synuclein within astrocytic puncta (Fig. 4M-P). This may suggest the initiation of neuroprotection against the incomplete lysosomal degradation of α-synuclein in the astrocytes. In the rat rotenone model, vitamin D treatment before and after the lesion partially restores SNC and striatal tyrosine hydroxylase expression [48, 49], and in three out of four clinical trials of vitamin D supplementation in PD, enhanced physical performance and reduced L-dopa dyskinesias were reported [50], suggesting that vitamin D can ameliorate cellular dysfunctions associated with α-synuclein aggregations. This function may be initiated in astrocytes at a later disease stage to that assessed in the present study.

Overall, we have shown that astrocytes in this A53T transgenic mouse model upregulate their clearance of mutant α-synuclein aggregates from the tissue analysed, suggesting this upregulation occurs earlier than any substantial neuronal accumulation. As indicated, the tissue changes are most consistent with astrocyte-dependent circuit remodelling by synapse phagocytosis. While the CP terminals most amenable to synaptic remodelling are likely to be dopaminergic terminals, the afferents to the SNC that might be involved could be either from the CP, prefrontal, cingulate, insular and/or somatomotor cortices [51] or from the retrorubral field that contains the A8 dopaminergic cell group [52]. The loss of nigral dopamine release [53] or the selective loss of A8 dopamine neurons [54] induces progressive parkinsonism in animal models, suggestive that the terminals most targeted for remodelling could be the dopaminergic terminals in all regions. Importantly, in early PD there is a loss of presynaptic terminals only in the SNC that accompanies the loss of dopamine function in synapses in the CP at the same time [55], a finding recently validated and shown to increase with disease progression [56]. While speculative, a loss of the ability of astrocytes to remodel synapses and mop up α-synuclein in these regions may be the precipitating event for the degeneration in PD.

## List of abbreviations

ALDH1L1: 10-formyltetrahydrofolate dehydrogenase
ROI: region of interest
BSA: bovine serum albumin
RRID: research resource identifier
CP: striatum
S100β: S100 calcium binding protein B
CYP27B1: 25-hydroxyvitamin D 1-a-hydroxylase
SNC: substantia nigra pars compacta
GFAP: glial fibrillary acidic protein
TBS: tris-buffered saline
PBS: phosphate buffered saline
VEGFA: vascular endothelial growth factor A
PD: Parkinson’s disease
VTA: ventral tegmental area

## Acknowledgements

The authors acknowledge: Amelia Sedjahtera for breeding, genotyping, and culling animals, and cutting the sections used in the analysis; Microscopy Australia at the Sydney Microscopy and Microanalysis facility, University of Sydney, enabled by NCRIS; and Heidi Cartwright for assistance with the figures.

## Authors contributions

Conceptualisation and study design: Cormac Peat, Asheeta Prasad and Glenda Halliday as part of a larger funded project designed by Carolyn Sue, Jennifer Johnston, Claire Parish, Lachlan Thompson, Deniz Kirik and Glenda Halliday. Reagent/materials/analysis tools: David Finkelstein, Asheeta Prasad and Glenda Halliday. Data acquisition: Cormac Peat. Data analysis and interpretation: Cormac Peat, Asheeta Prasad and Glenda Halliday. Drafting of the manuscript: Cormac Peat and Glenda Halliday. Editing of the manuscript: All authors. All authors have read and approved the final manuscript

## Data availability

The data, code, protocols, and key lab materials used and generated in this study are listed in a Key Resource Table alongside their persistent identifiers at Zenodo https://zenodo.org/records/18345432. Images analysed are deposited at https://www.ebi.ac.uk/biostudies/studies/S-BIAD2485 and generated data and readme files are deposited at Zenodo https://zenodo.org/records/18345432.

## Ethical approval

All animal procedures were conducted in agreement with the Australian National Health and Medical Research Council’s published Code of Practice for the Use of Animals in Research, and approval granted by The Florey Institute of Neuroscience and Mental Health Animal Ethics committee (22-034-FINMH).

## References

1 Poewe W, Seppi K, Tanner CM, Halliday GM, Brundin P, Volkmann J, Schrag AE, Lang AE. Parkinson disease. Nat Rev Dis Primers 2017; 3: 17013

2 Simuni T, Chahine LM, Poston K, Brumm M, Buracchio T, Campbell M, Chowdhury S, Coffey C, Concha-Marambio L, Dam T, DiBiaso P, Foroud T, Frasier M, Gochanour C, Jennings D, Kieburtz K, Kopil CM, Merchant K, Mollenhauer B, Montine T, Nudelman K, Pagano G, Seibyl J, Sherer T, Singleton A, Stephenson D, Stern M, Soto C, Tanner CM, Tolosa E, Weintraub D, Xiao Y, Siderowf A, Dunn B, Marek K. A biological definition of neuronal alpha-synuclein disease: towards an integrated staging system for research. Lancet Neurol 2024; 23: 178–90

3 Braak H, Sastre M, Del Tredici K. Development of alpha-synuclein immunoreactive astrocytes in the forebrain parallels stages of intraneuronal pathology in sporadic Parkinson’s disease. Acta Neuropathol 2007; 114: 231–41

4 Kovacs GG, Breydo L, Green R, Kis V, Puska G, Lorincz P, Perju-Dumbrava L, Giera R, Pirker W, Lutz M, Lachmann I, Budka H, Uversky VN, Molnar K, Laszlo L. Intracellular processing of disease-associated alpha-synuclein in the human brain suggests prion-like cell-to-cell spread. Neurobiol Dis 2014; 69: 76–92

5 Wakabayashi K, Hayashi S, Yoshimoto M, Kudo H, Takahashi H. NACP/alpha-synuclein-positive filamentous inclusions in astrocytes and oligodendrocytes of Parkinson’s disease brains. Acta Neuropathol 2000; 99: 14–20

6 Huynh B, Fu Y, Kirik D, Shine JM, Halliday GM. Comparison of Locus Coeruleus Pathology with Nigral and Forebrain Pathology in Parkinson’s Disease. Mov Disord 2021; 36: 2085–93

7 Song YJ, Halliday GM, Holton JL, Lashley T, O’Sullivan SS, McCann H, Lees AJ, Ozawa T, Williams DR, Lockhart PJ, Revesz TR. Degeneration in different parkinsonian syndromes relates to astrocyte type and astrocyte protein expression. J Neuropathol Exp Neurol 2009; 68: 1073–83

8 Altay MF, Liu AKL, Holton JL, Parkkinen L, Lashuel HA. Prominent astrocytic alpha-synuclein pathology with unique post-translational modification signatures unveiled across Lewy body disorders. Acta Neuropathol Commun 2022; 10: 163

9 Loria F, Vargas JY, Bousset L, Syan S, Salles A, Melki R, Zurzolo C. alpha-Synuclein transfer between neurons and astrocytes indicates that astrocytes play a role in degradation rather than in spreading. Acta Neuropathol 2017; 134: 789–808

10 Halliday GM, Stevens CH. Glia: initiators and progressors of pathology in Parkinson’s disease. Mov Disord 2011; 26: 6–17

11 Mirza B, Hadberg H, Thomsen P, Moos T. The absence of reactive astrocytosis is indicative of a unique inflammatory process in Parkinson’s disease. Neuroscience 2000; 95: 425–32

12 Tong J, Ang LC, Williams B, Furukawa Y, Fitzmaurice P, Guttman M, Boileau I, Hornykiewicz O, Kish SJ. Low levels of astroglial markers in Parkinson’s disease: relationship to alpha-synuclein accumulation. Neurobiol Dis 2015; 82: 243–53

13 Ma M, Paryani F, Jakubiak K, Xia S, Antoku S, Kannan A, Lee J, Madden N, Senthil Kumar S, Li J, Chen D, Hargus G, Mahajan A, Flowers X, Harms AS, Sulzer D, Goldman JE, Sims PA, Al-Dalahmah O. The spatial landscape of glial pathology and T cell response in Parkinson’s disease substantia nigra. Nat Commun 2025; 16: 7146

14 Mazzetti S, Barichella M, Giampietro F, Giana A, Calogero AM, Amadeo A, Palazzi N, Comincini A, Giaccone G, Bramerio M, Caronni S, Cereda V, Cereda E, Cappelletti G, Rolando C, Pezzoli G. Astrocytes expressing Vitamin D-activating enzyme identify Parkinson’s disease. CNS Neurosci Ther 2022; 28: 703–13

15 Giasson BI, Duda JE, Quinn SM, Zhang B, Trojanowski JQ, Lee VM. Neuronal alpha-synucleinopathy with severe movement disorder in mice expressing A53T human alphasynuclein. Neuron 2002; 34: 521–33

16 Zhang TD, Kolbe SC, Beauchamp LC, Woodbridge EK, Finkelstein DI, Burrows EL. How Well Do Rodent Models of Parkinson’s Disease Recapitulate Early Non-Motor Phenotypes? A Systematic Review. Biomedicines 2022; 10:

17 Ke M, Chong CM, Zhu Q, Zhang K, Cai CZ, Lu JH, Qin D, Su H. Comprehensive Perspectives on Experimental Models for Parkinson’s Disease. Aging Dis 2021; 12: 223–46

18 Gu XL, Long CX, Sun L, Xie C, Lin X, Cai H. Astrocytic expression of Parkinson’s disease-related A53T alpha-synuclein causes neurodegeneration in mice. Mol Brain 2010; 3: 12

19 Lan G, Wang P, Chan RB, Liu Z, Yu Z, Liu X, Yang Y, Zhang J. Astrocytic VEGFA: An essential mediator in blood-brain-barrier disruption in Parkinson’s disease. Glia 2022; 70: 337–53

20 Hinkle JT, Dawson VL, Dawson TM. The A1 astrocyte paradigm: New avenues for pharmacological intervention in neurodegeneration. Mov Disord 2019; 34: 959–69

21 Escartin C, Galea E, Lakatos A, O’Callaghan JP, Petzold GC, Serrano-Pozo A, Steinhauser C, Volterra A, Carmignoto G, Agarwal A, Allen NJ, Araque A, Barbeito L, Barzilai A, Bergles DE, Bonvento G, Butt AM, Chen WT, Cohen-Salmon M, Cunningham C, Deneen B, De Strooper B, Diaz-Castro B, Farina C, Freeman M, Gallo V, Goldman JE, Goldman SA, Gotz M, Gutierrez A, Haydon PG, Heiland DH, Hol EM, Holt MG, Iino M, Kastanenka KV, Kettenmann H, Khakh BS, Koizumi S, Lee CJ, Liddelow SA, MacVicar BA, Magistretti P, Messing A, Mishra A, Molofsky AV, Murai KK, Norris CM, Okada S, Oliet SHR, Oliveira JF, Panatier A, Parpura V, Pekna M, Pekny M, Pellerin L, Perea G, Perez-Nievas BG, Pfrieger FW, Poskanzer KE, Quintana FJ, Ransohoff RM, Riquelme-Perez M, Robel S, Rose CR, Rothstein JD, Rouach N, Rowitch DH, Semyanov A, Sirko S, Sontheimer H, Swanson RA, Vitorica J, Wanner IB, Wood LB, Wu J, Zheng B, Zimmer ER, Zorec R, Sofroniew MV, Verkhratsky A. Reactive astrocyte nomenclature, definitions, and future directions. Nat Neurosci 2021; 24: 312–25

22 Li J, Pan L, Pembroke WG, Rexach JE, Godoy MI, Condro MC, Alvarado AG, Harteni M, Chen YW, Stiles L, Chen AY, Wanner IB, Yang X, Goldman SA, Geschwind DH, Kornblum HI, Zhang Y. Conservation and divergence of vulnerability and responses to stressors between human and mouse astrocytes. Nat Commun 2021; 12: 3958

23 Qian Z, Qin J, Lai Y, Zhang C, Zhang X. Large-Scale Integration of Single-Cell RNA-Seq Data Reveals Astrocyte Diversity and Transcriptomic Modules across Six Central Nervous System Disorders. Biomolecules 2023; 13:

24 Tatsumi K, Isonishi A, Yamasaki M, Kawabe Y, Morita-Takemura S, Nakahara K, Terada Y, Shinjo T, Okuda H, Tanaka T, Wanaka A. Olig2-Lineage Astrocytes: A Distinct Subtype of Astrocytes That Differs from GFAP Astrocytes. Front Neuroanat 2018; 12: 8

25 Linnerbauer M, Wheeler MA, Quintana FJ. Astrocyte Crosstalk in CNS Inflammation. Neuron 2020; 108: 608–22

26 Zhong J, Tang G, Zhu J, Wu W, Li G, Lin X, Liang L, Chai C, Zeng Y, Wang F, Luo L, Li J, Chen F, Huang Z, Zhang X, Zhang Y, Liu H, Qiu X, Tang S, Chen D. Single-cell brain atlas of Parkinson’s disease mouse model. J Genet Genomics 2021; 48: 277–88

27 Diwakarla S, McQuade RM, Constable R, Artaiz O, Lei E, Barnham KJ, Adlard PA, Cherny RA, Di Natale MR, Wu H, Chai XY, Lawson VA, Finkelstein DI, Furness JB. ATH434 Reverses Colorectal Dysfunction in the A53T Mouse Model of Parkinson’s Disease. J Parkinsons Dis 2021; 11: 1821–32

28 Jaisa-Aad M, Munoz-Castro C, Healey MA, Hyman BT, Serrano-Pozo A. Characterization of monoamine oxidase-B (MAO-B) as a biomarker of reactive astrogliosis in Alzheimer’s disease and related dementias. Acta Neuropathol 2024; 147: 66

29 Low CYB, Lee JH, Lim FTW, Lee C, Ballard C, Francis PT, Lai MKP, Tan MGK. Isoform-specific upregulation of FynT kinase expression is associated with tauopathy and glial activation in Alzheimer’s disease and Lewy body dementias. Brain Pathol 2021; 31: 253–66

30 Yang Y, Song JJ, Choi YR, Kim SH, Seok MJ, Wulansari N, Darsono WHW, Kwon OC, Chang MY, Park SM, Lee SH. Therapeutic functions of astrocytes to treat alpha-synuclein pathology in Parkinson’s disease. Proc Natl Acad Sci U S A 2022; 119: e2110746119

31 Lindstrom V, Gustafsson G, Sanders LH, Howlett EH, Sigvardson J, Kasrayan A, Ingelsson M, Bergstrom J, Erlandsson A. Extensive uptake of alpha-synuclein oligomers in astrocytes results in sustained intracellular deposits and mitochondrial damage. Mol Cell Neurosci 2017; 82: 143–56

32 Chavarria C, Rodriguez-Bottero S, Quijano C, Cassina P, Souza JM. Impact of monomeric, oligomeric and fibrillar alpha-synuclein on astrocyte reactivity and toxicity to neurons. Biochem J 2018; 475: 3153–69

33 Tsunemi T, Ishiguro Y, Yoroisaka A, Valdez C, Miyamoto K, Ishikawa K, Saiki S, Akamatsu W, Hattori N, Krainc D. Astrocytes Protect Human Dopaminergic Neurons from alpha-Synuclein Accumulation and Propagation. J Neurosci 2020; 40: 8618–28

34 Muwanigwa MN, Modamio-Chamarro J, Antony PMA, Gomez-Giro G, Kruger R, Bolognin S, Schwamborn JC. Alpha-synuclein pathology is associated with astrocyte senescence in a midbrain organoid model of familial Parkinson’s disease. Mol Cell Neurosci 2024; 128: 103919

35 Simmnacher K, Krach F, Schneider Y, Alecu JE, Mautner L, Klein P, Roybon L, Prots I, Xiang W, Winner B. Unique signatures of stress-induced senescent human astrocytes. Exp Neurol 2020; 334: 113466

36 Knott C, Wilkin GP, Stern G. Astrocytes and microglia in the substantia nigra and caudate-putamen in Parkinson’s disease. Parkinsonism Relat Disord 1999; 5: 115–22

37 Park J, Chung WS. Astrocyte-dependent circuit remodeling by synapse phagocytosis. Curr Opin Neurobiol 2023; 81: 102732

38 Shen W, Zhai S, Surmeier DJ. Striatal synaptic adaptations in Parkinson’s disease. Neurobiol Dis 2022; 167: 105686

39 Morales I, Sanchez A, Rodriguez-Sabate C, Rodriguez M. Striatal astrocytes engulf dopaminergic debris in Parkinson’s disease: A study in an animal model. PLoS One 2017; 12: e0185989

40 Sun L, Zheng X, Che Y, Zhang Y, Huang Z, Jia L, Zhu Y, Lei W, Guo G, Shao C. Morphological changes in perisynaptic astrocytes induced by dopamine neuronal degeneration in the striatum of rats. Heliyon 2024; 10: e27637

41 Zhu YF, Wang WP, Zheng XF, Chen Z, Chen T, Huang ZY, Jia LJ, Lei WL. Characteristic response of striatal astrocytes to dopamine depletion. Neural Regen Res 2020; 15: 724–30

42 Izco M, Blesa J, Verona G, Cooper JM, Alvarez-Erviti L. Glial activation precedes alpha-synuclein pathology in a mouse model of Parkinson’s disease. Neurosci Res 2021; 170: 330–40

43 Elabi O, Gaceb A, Carlsson R, Padel T, Soylu-Kucharz R, Cortijo I, Li W, Li JY, Paul G. Human alpha-synuclein overexpression in a mouse model of Parkinson’s disease leads to vascular pathology, blood brain barrier leakage and pericyte activation. Sci Rep 2021; 11: 1120

44 Wada K, Arai H, Takanashi M, Fukae J, Oizumi H, Yasuda T, Mizuno Y, Mochizuki H. Expression levels of vascular endothelial growth factor and its receptors in Parkinson’s disease. Neuroreport 2006; 17: 705–9

45 Faucheux BA, Bonnet AM, Agid Y, Hirsch EC. Blood vessels change in the mesencephalon of patients with Parkinson’s disease. Lancet 1999; 353: 981–2

46 Elgayar SAM, Abdel-Hafez AAM, Gomaa AMS, Elsherif R. Vulnerability of glia and vessels of rat substantia nigra in rotenone Parkinson model. Ultrastruct Pathol 2018; 42: 181–92

47 Yang P, Pavlovic D, Waldvogel H, Dragunow M, Synek B, Turner C, Faull R, Guan J. String Vessel Formation is Increased in the Brain of Parkinson Disease. J Parkinsons Dis 2015; 5: 821–36

48 Sanchez B, Relova JL, Gallego R, Ben-Batalla I, Perez-Fernandez R. 1,25-Dihydroxyvitamin D3 administration to 6-hydroxydopamine-lesioned rats increases glial cell line-derived neurotrophic factor and partially restores tyrosine hydroxylase expression in substantia nigra and striatum. J Neurosci Res 2009; 87: 723–32

49 Lima LAR, Lopes MJP, Costa RO, Lima FAV, Neves KRT, Calou IBF, Andrade GM, Viana GSB. Vitamin D protects dopaminergic neurons against neuroinflammation and oxidative stress in hemiparkinsonian rats. J Neuroinflammation 2018; 15: 249

50 Homann CN, Homann B, Ivanic G, Urbanic-Purkart T. Vitamin D supplementation in later life: a systematic review of efficacy and safety in movement disorders. Front Aging Neurosci 2024; 16: 1333217

51 Zhang Y, Larcher KM, Misic B, Dagher A. Anatomical and functional organization of the human substantia nigra and its connections. Elife 2017; 6:

52 Arts MP, Groenewegen HJ, Veening JG, Cools AR. Efferent projections of the retrorubral nucleus to the substantia nigra and ventral tegmental area in cats as shown by anterograde tracing. Brain Res Bull 1996; 40: 219–28

53 Gonzalez-Rodriguez P, Zampese E, Stout KA, Guzman JN, Ilijic E, Yang B, Tkatch T, Stavarache MA, Wokosin DL, Gao L, Kaplitt MG, Lopez-Barneo J, Schumacker PT, Surmeier DJ. Disruption of mitochondrial complex I induces progressive parkinsonism. Nature 2021; 599: 650–6

54 Deutch AY, Elsworth JD, Goldstein M, Fuxe K, Redmond DE, Jr., Sladek JR, Jr., Roth RH. Preferential vulnerability of A8 dopamine neurons in the primate to the neurotoxin 1-methyl-4-phenyl-1,2,3,6-tetrahydropyridine. Neurosci Lett 1986; 68: 51–6

55 Delva A, Van Weehaeghe D, Koole M, Van Laere K, Vandenberghe W. Loss of Presynaptic Terminal Integrity in the Substantia Nigra in Early Parkinson’s Disease. Mov Disord 2020; 35: 1977–86

56 Holmes SE, Honhar P, Tinaz S, Naganawa M, Hilmer AT, Gallezot JD, Dias M, Yang Y, Toyonaga T, Esterlis I, Mecca A, Van Dyck C, Henry S, Ropchan J, Nabulsi N, Louis ED, Comley R, Finnema SJ, Carson RE, Matuskey D. Synaptic loss and its association with symptom severity in Parkinson’s disease. NPJ Parkinsons Dis 2024; 10: 42

